# Tissue-specific accumulation profiles of phorbol esters in *Jatropha curcas* and gene induction in response to abiotic and biotic stresses

**DOI:** 10.1101/2020.09.26.315010

**Authors:** Lei Wei, Wei Zhang, Caixin Fan, Tingwei Dai, Shijuan Li, Fang Chen, Ying Xu

**Affiliations:** Key Laboratory of Bio-resources and Eco-environment, Ministry of Education, College of Life Sciences, Sichuan University, Chengdu, 29 Wangjiang Road, 610064, China; Biotechnology Developing Center, Henan Academy of Sciences, Zhengzhou, Hongzhuan Road, 450002, China; Department of Plant Pathology, Kansas State University, 1712 Claflin Road, Throckmorton Hall, Manhattan, KS, 66506, USA; Key Laboratory of Resource Biology and Biopharmaceutical Engineering, College of Life Sciences, Sichuan University, Chengdu, 29 Wangjiang Road, 610064, China

**Keywords:** Phorbol esters, *Jatropha curcas*, Terpenoids, Gene expression, Pathway, Stress

## Abstract

*Jatropha curcas* L. (*J. curcas*), a shrub plant of the *Euphorbiaceae* family, has received enormous attention as a promising biofuel plant for the production of biodiesel and medical potential in ethnopharmacology. However, the tumor-promoter toxin phorbol esters present in *J. curcas* raises concerns for health and environmental risk as its large-scale cultivation limits the use of meal obtained after oil extraction for animal feed. Here, we determined the variation of phorbol ester profiles and contents in eight *J. curcas* tissues by high performance liquid chromatography (HPLC) and found phorbol esters present in all parts of the plant except the seed shell. We showed tissue-specific patterns of accumulation of phorbol esters and associated terpenoids at the transcriptomic level with high transcript levels in reproductive and young tissues. Genes involved in the same module of terpenoids biosynthesis were positively correlated. We further present diverse abiotic and biotic stresses that had different effects on the accumulation of transcripts in terpenoids shared and branched terpenoid pathways in plant seedlings. The fine-tuning of terpenoids biosynthesis may link with ecological functions in plants under extreme environments and defense against pathogens.

## 1.1. Introduction

*Jatropha curcas* (*J. curcas, Euphorbiaceae*) is a multipurpose shrub distributed widely in tropical and subtropical regions of Latin America, Asia, and Africa (Montes and Melchinger, 2016). Its ability to survive under extreme conditions, such as heat and drought, makes it popular as domesticated hedge in barren lands (Krishnamurthy et al., 2012). Recently, *J. curcas* has gained enormous attention as a promising feedstock crop in the biodiesel industry (Berchmans and Hirata, 2008; Devappa et al., 2010a). *J. curcas* seeds contain oil that can be used as a diesel substitute in the biodiesel industry. Besides, seed meals, the co-product after oil extraction, contain high-level proteins and can serve as good sources for livestock (King et al., 2009; Makkar et al., 2008). However, the presence of toxic phorbol esters in seeds and other parts of *J. curcas* dramatically restricts the widespread use of its oil and seed meal because the tumor-promoting activity of phorbol esters may pose risks on human health and the environment (Hirota et al., 1988; Makkar et al., 1997; Saetae and Suntornsuk, 2011).

Presently, six types of phorbol esters have been isolated from *J. curcas* seed oil (Haas et al., 2002). These esters share the same phorbol core, a tetracyclic carbon skeleton, with diverse intramolecular di-esters. Although accumulated as a phytoalexin in plants to facilitate their fitness, phorbol esters are known for their extensive bioactivities, especially as a carcinogen, in a wide range of organisms. For example, they can induce acute toxic effects in animals and insets and cause skin irritant and promote tumors in human beings (Becker and Makkar, 1998; Devappa et al., 2012; Katole et al., 2011; Li et al., 2010; Rug and Ruppel, 2000). In contrast, recent studies revealed phorbol esters can exhibit antimicrobial and antitumor activities when applied at low concentrations (Devappa et al., 2013b; Fujiki et al., 2017). The diverse bioactivities of phorbol esters are not only dependent on dosage effects but also associated with varied structures among phorbol esters and their derivates. These varied structures are the basis for phorbol esters to activate protein kinase C involved signaling transduction, which is involved in diverse cellular processes and promotes tumors. Indeed, the ethnobotanical use of *J. curcas* is not only attributed to some pharmaceutical compounds but also associated with toxins, such as phorbol esters and protein curcin (Debnath and Bisen, 2008; Devappa et al., 2010b; Igbinosa et al., 2011; Insanu et al., 2013; Joseph, 2001; KOSASI et al., 1989; Lin et al., 2003; Qin et al., 2005; Ravindranath et al., 2004; Sabandar et al., 2013; Wei et al., 2015). Elucidating the types and abundance of phorbol esters in different parts of *J. curcas* is necessary for harnessing this toxic plant as a value to the biodiesel industry and with high pharmaceutical potential.

Phorbol esters are diterpenoids that are synthesized via the terpenoid pathway mainly in *Euphorbiaceae* and *Thymelaeaceae* family. Their biosynthesis can be divided into three separate phases: a) formation of the phorbol core, the C_20_ tetracyclic skeleton, b) esterification of polyhydroxy groups at C-12, C-13, and C-20 positions of the phorbol core, and c) further modification of the carbon skeleton and side chains (King et al., 2014). In plants, formation of C_20_ diterpene skeleton precursor gernanylgeranyl diphosphate (GGPP) is catalyzed by gernanylgeranyl diphosphate synthase (GGPPs) and requires the two C_5_ isoprenoid precursor, isopentenyl diphosphate (IPP), and its isomer dimethylallyl diphosphate (DMAPP, Lange et al., 2000). These precursors derived from mevalonate (MVA) pathway or non-MVA pathway (MEP or DXP pathway) are required by all terpenoid productions, including geranyl diphosphate (GPP, C_10_) derived monoterpenoids, farnesyl diphosphate (FPP, C_15_) derived sesquiterpenes and triterpenoids, and geranylgeranyl diphosphate (GGPP, C_20_) derived diterpenoids (Tholl, 2015; Vranová et al., 2013). The GGPP backbones can be further cyclized to form the phorbol core by a terpene synthase named casbene synthase (CAS) (Li et al., 2016; Nakano et al., 2012). Then, some modification enzymes, such as acyltransferase, terpene hydroxylase, and cytochrome P_450_monooxygenases and reductases, are responsible for adding ester groups to side chains and further modification (Zerbe and Bohlmann, 2015). Terpenoid accumulation displays tissue- and developmental stage-specific patterns and can be induced upon abiotic and biotic stresses (Pichersky and Raguso, 2018; Yazaki et al., 2017). Although *Euphorbiaceae* plants are tolerant of heat and drought, little is known about how they manipulate terpenoid production, especially the major phytoalexin phorbol esters, to adapt to abiotic and biotic stresses.

In this study, we investigated the accumulation of phorbol esters and associated gene expression across different *J. curcas* tissues. Using HPLC, we show that phorbol esters are found in all tested *J. curcas* tissues except seed shells at varying types and abundances. The quantitative RT-PCR revealed that the expression profiles of genes involved in phorbol esters and terpenoids biosynthesis exhibited tissue-specific patterns with higher accumulations in reproductive organs and young tissues. Besides, we found that genes involved in terpenoids production in *J. curcas* are differentially regulated in response to abiotic and biotic stressors. In summary, phorbol ester content and gene expression data provide support for the reduction of phorbol ester content through chemical detoxification or breeding and genome editing. Such detoxification are required for large-scale application of *J. curcas* oil in the biofuel industry, seed meal in agriculture, and bioactive components in pharmaceuticals. The regulation of terpenoid gene expression in response to abiotic and biotic stresses such as drought and fungal pathogens indicates that terpenoids, especially phorbol esters, play an important role in *Euphorbiaceae* and *Thymelaeaceae* plants’ tolerance against the extreme environmental condition.

## 2. Results

### 2.1. Phorbol ester metabolic profiling show a tissue-specific pattern

To investigate the types and abundance of phorbol esters in *J. curcas*, we determined the metabolic profile of phorbol esters in different plant parts. We first established the standard by determination of phorbol ester profile in seeds by HPLC based on previous studies (Devappa et al., 2013a; Hua et al., 2015; Liu et al., 2013; Wang et al., 2012). We identified five phorbol esters peaks containing phorbol ester factors (Jc1, Jc2, Jc3, Jc4 & Jc5, and Jc6) based on their retention times (Figure 1A). To effectively and consistently extract phorbol esters from different parts of *J. curcas*, we developed a sonication-assisted extraction method and optimized five extraction parameters, including solvent, soaking temperature, solvent to material ratio, extraction time, and sonication temperature. Analysis of variance (ANOVA) revealed that the extraction solvent had the most significant impacts on phorbol esters yield (Supplemental Table 1). The highest yield was obtained at the optimized parameters (Supplemental Figure 1). Thereafter, we used the optimized extraction method coupled with HPLC determination for all phorbol ester content analysis in this study.

**Table 1.**
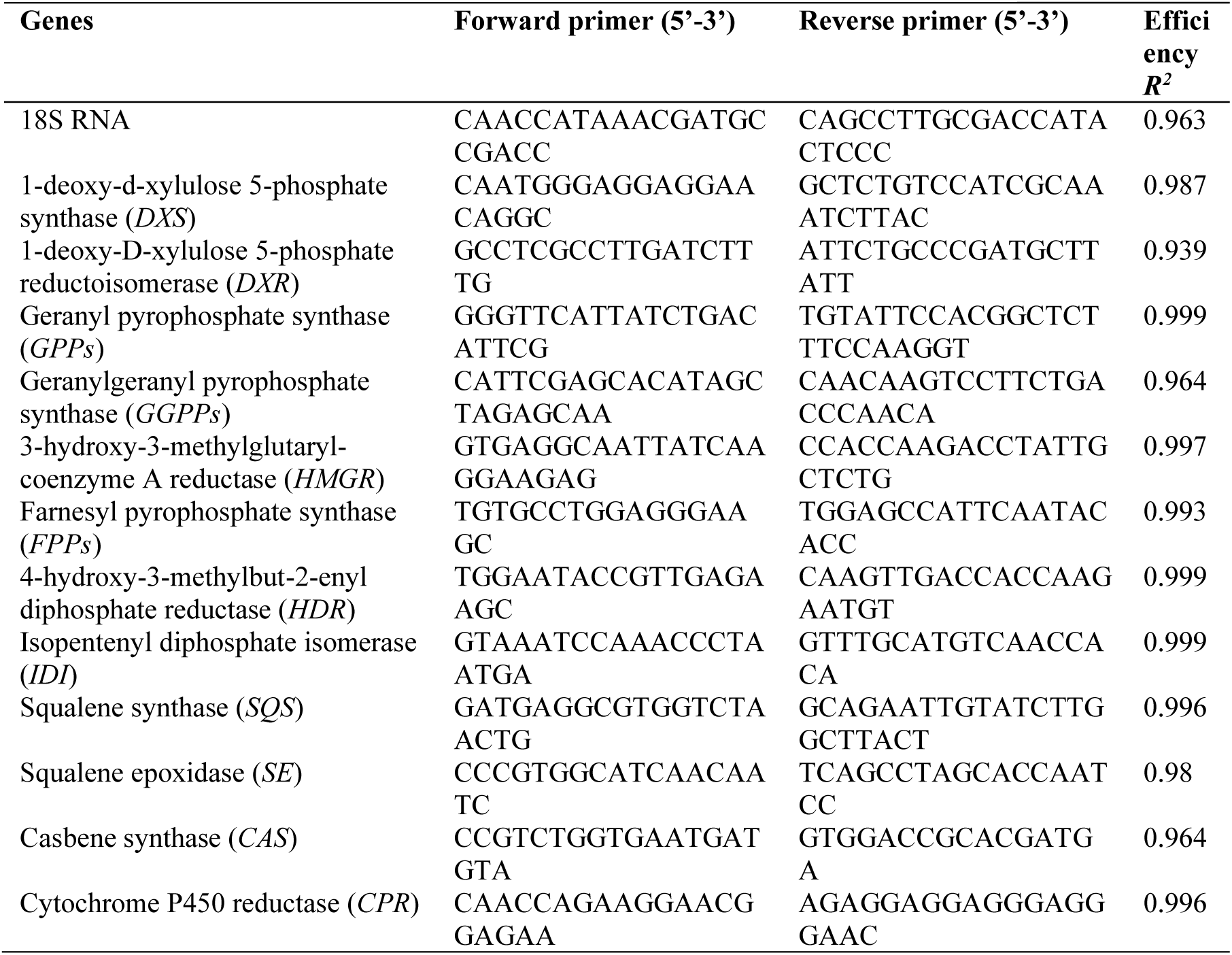
Nucleotide sequences and efficiency of primers used in qPCR.

**Figure 1.**
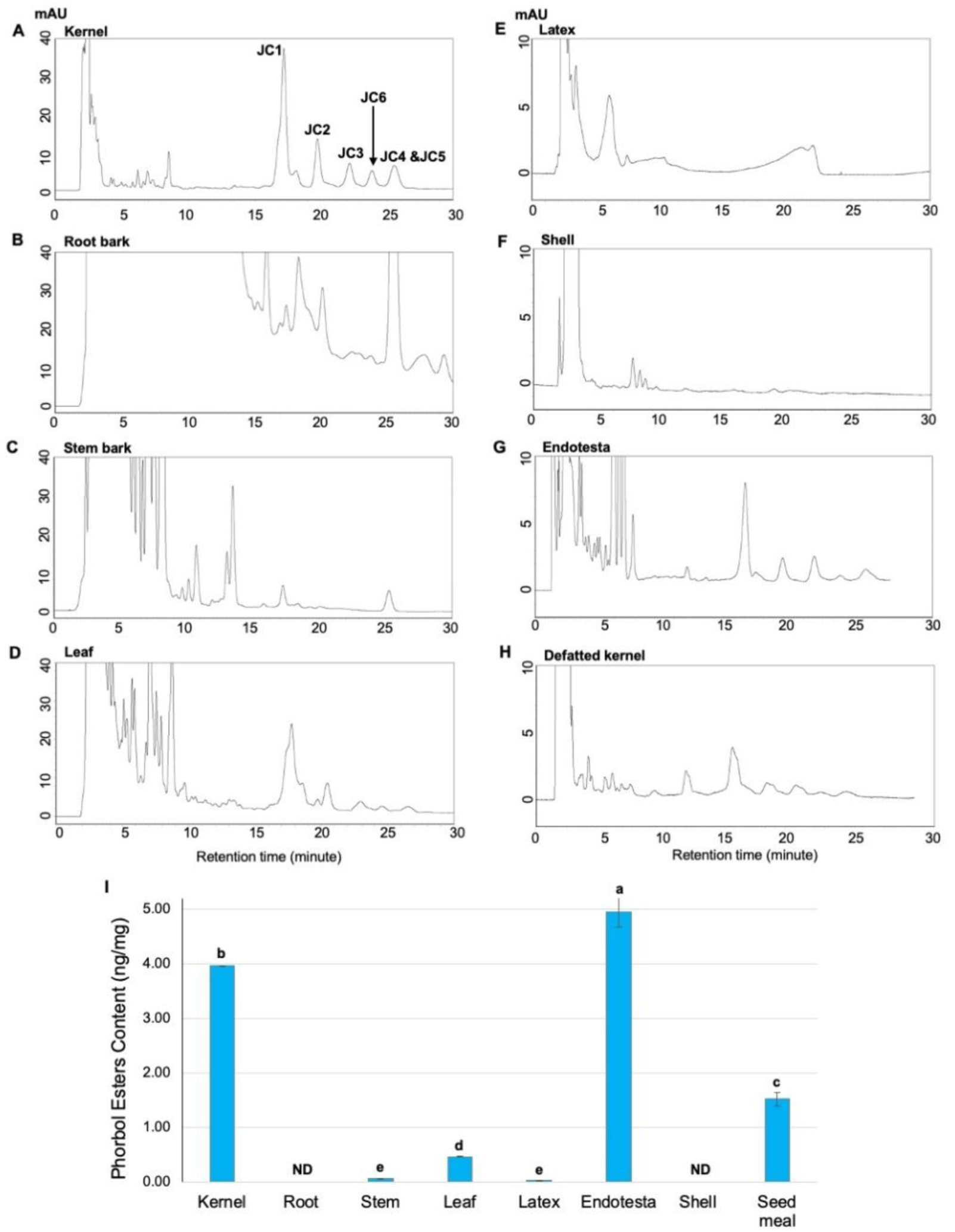
HPLC analysis of phorbol esters in *Jatropha curcas*. HPLC profiles of six types of phorbol esters extracted from (A) kernel, (B) root, (C) young stem, (D) wild leaf, (E) latex, (F) endotesta, (G) shell, and (H) seed meal. PMA was used as external standards. Jatropha factors 1 to 6 are labeled. (I) Total contents of six phorbol esters in diverse *J. curcas* tissues were determined by calibrating with a standard curve of PMA. The data represent means ± standard error from triplicate biological replicates. ND = not detected. Letters at the top of 2I indicate statistically homogeneours groups determined by HSD test.

We identified varied types and abundance of phorbol esters in root bark, stem bark, leaf, latex, shell, and endotesta by comparing the retention time with kernel extraction (Figure 1). We identified all six phorbol esters factors, Jc1 to Jc6, from five plant tissues and seed meals except root bark. The extraction and determination methods used in this study failed to identify phorbol ester in root barks due to the presence of impurities at the background at the same retention time (Figure 1B). Comparison of the chromatogram profiles obtained from different tissues revealed that both types and abundance of phorbol esters varied among plant tissues. For example, endotesta and leaves shared a similar phorbol ester composition as kernels but varied at total content. Whereas, latex and stem bark showed only one or two Jc factors compared with the complete phorbol ester profile in kernels (Figure 1C and 1E). The endotesta showed the highest level of total phorbol esters among all examined tissues (Figure 1G and 1F). Leaves, latex, and stem bark, which are usually used as traditional medicines, showed very low amounts of phorbol esters. The seed meal showed dramatically reduced but still considerable content of phorbol esters (1.517+-0.120 mg/g, Figure 2I), suggesting that detoxification is required before use as a feed resource. No detectable phorbol esters were identified in seed shells.

**Figure 2.**
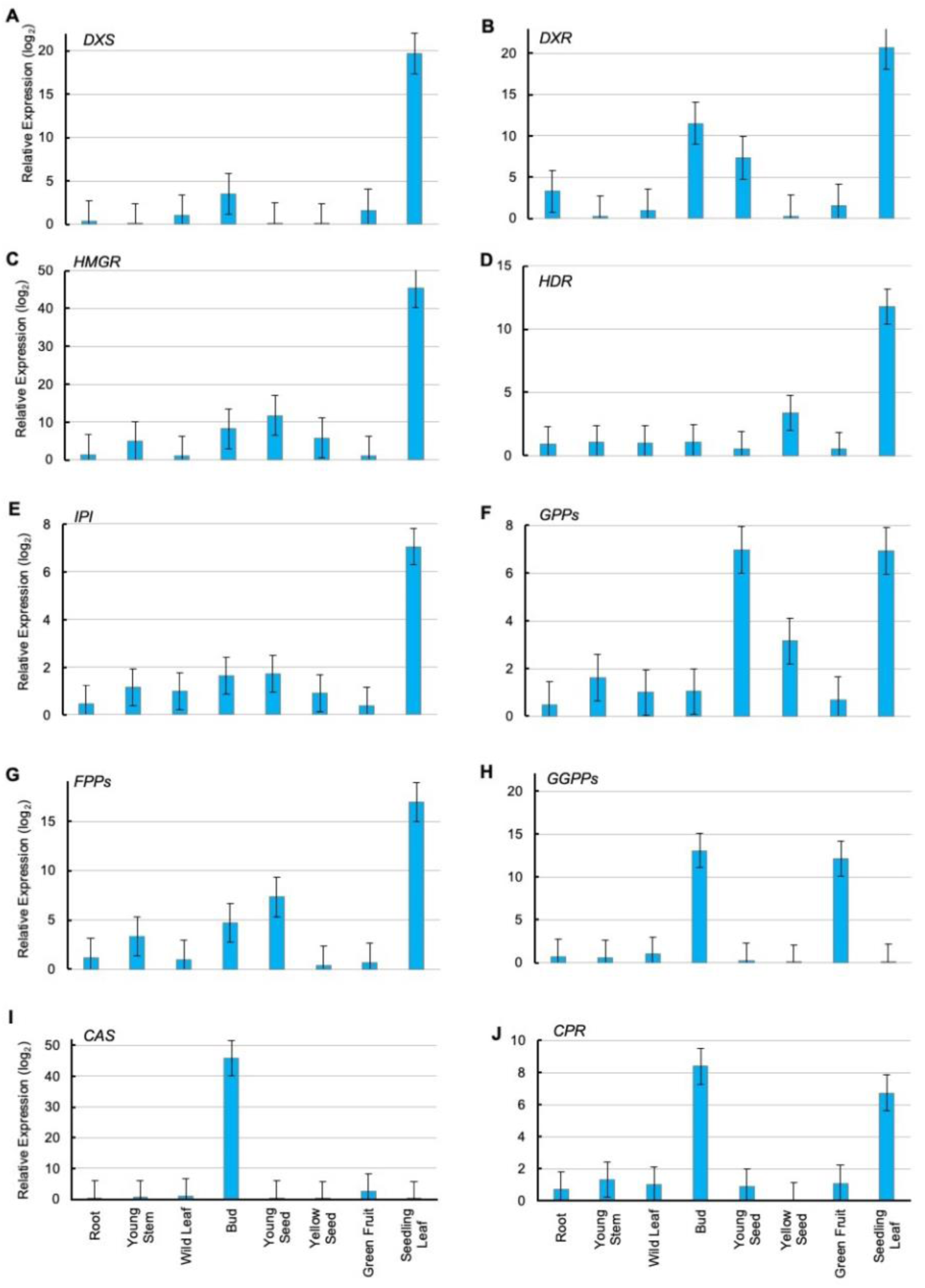
Expression of genes involved in terpene metabolism and phorbol esters biosynthesis pathway in different tissues of *Jatropha curcas*. Accumulation levels of transcripts were determined by qRT-PCR using primers in Table 1. Relative expression levels of individual transcripts in diverse tissues were calculated by comparison of field-growing mature leaf, including root, young stem, bud, young seed, yellow seed, green fruit, seedling leaf. All graphs represent the values of three replicates with cDNA prepared from different tissues. Each bar represents means ± standard error.

### 2.2. Phorbol esters prefer to accumulate in plant reproductive and young tissues

To understand the molecular mechanisms controlling phorbol ester tissue-specific accumulation, we investigated phorbol ester associated gene expression profile across eight plant tissues using quantitative reverse transcription polymerase chain reaction (qRT-PCR, Table 1 and Figure 2). All gene expression profiles from *J. curcas* tissues are standardized to their expression in field-grown wild leaf tissue. We targeted phorbol esters’ pathway-specific genes, including *GGPPs* encoding enzyme for diterpene precursor GGPP, *CAS* responsible for cyclization of phorbol core. Both are highly expressed in buds (13.1 and 45.9 fold change) and green fruit (2.7 and 12.14 fold change, Figure 2H and 2I). A cytochrome P_450_ reductase (*CPR*) responsible for secondary modification of phorbol core and side chains also showed a relatively high expression in flower buds (8.37-fold change) and green fruits (6.72-fold change) (Figure 2J). The up-regulation of phorbol ester pathway-specific genes in buds and green fruits suggests the biosynthesis of phorbol esters occurs at a higher rate in reproductive tissues.

We next analyzed the genes involved in both shared and branched isoprenoid pathways. Five genes involved in the shared pathway are responsible for C_5_ isoprenoid production, including *DXS* and *DXR* in the MEP pathway, *HMGR* and *HDR* in the MVA pathway, and *IPI* responsible for cytosolic IPP and DMAPP formation. We selected two genes, *GPPs* and *FPPs*, from branched pathways that essential for monoterpene, sesquiterpenoid, and triterpenoid production. The seven genes involved in isoprenoid shared and branched pathways showed the highest expression level in seedling leaves (Figure 2A-2E). For example, *DXS, DXR*, and *HMGR* showed remarkably high expression levels in seedling leaves with 19.65, 20.69, and 45.33 fold change. Four genes involved in both shared and branched pathways (*DXR, HMGR, GPPs*, and *FPPs*) also showed relatively higher expression levels in reproduction tissues of flower buds and young seeds (Figure 2B, 2C, 2F, and 2G). Collectively, phorbol ester genes were highly expressed in young and reproductive tissues compared with genes involved in common and other branch pathways, suggesting the protective roles of phorbol esters during plant development.

### 2.3. Terpenoids pathways showed a tissue-specific pattern

To gain insight into the relationship among diverse terpenoid pathways and their disproportionate accumulation among plant tissues, we tested correlations among genes condensed in different pathways based on their expression profile across eight tissues (Figure 3A). We identified four gene co-expression modules with genes showing strong positive correlations (Figure 3A). We also found that the four modules showed tissue-specific patterns, which were consistent with their biological functions. For instance, phorbol ester pathway-specific genes of *GGPPs* and *CAS* clustered in the same module with a strong positive correlation (R^2^ = 0.905, *P* = 0.005). Furthermore, they both are closely linked with genes from the MEP pathway but negatively correlated with the module containing genes from the cytosolic MVA pathway, indicating phorbol esters may derive from the MEP pathway in the plastid of *J. curcas* (Figure 3A). Whereas, the *GPPs* essential for monoterpene production is clustered together with *HMGR* and *IPI* in the MVA module, suggesting the C_10_ geranyl diphosphate (GDP)-based monoterpenoids may form through the cytosolic MVA pathway. The negative correlation between the diterpene module and the MVA module indicates a competition relationship between phorbol ester production and monoterpenoid production in *J. curcas*. Surprisingly, the *FPPs*, which is responsible for C15 sesquiterpene and C_30_ triterpene and steroid (two FDP units) production, also showed a closer relationship with genes in the MEP pathway.

**Figure 3.**
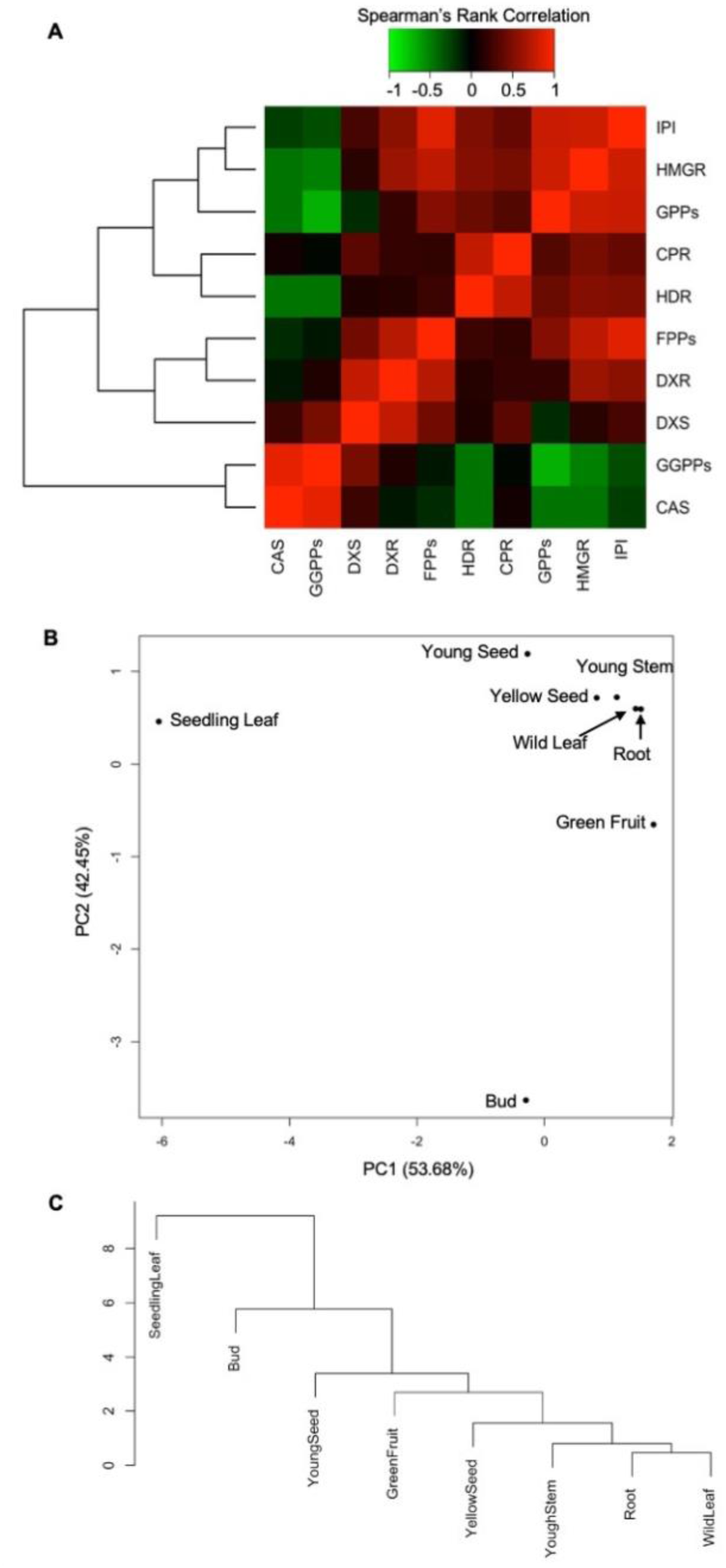
Tissue-specific patterns of transcripts accumulation involved in terpene metabolism and phorbol esters pathway in *Jatropha curcas*. (A) Heat map illustrates the Spearman’s rank correlation co-efficiency on expression levels of target genes involved terpene metabolism and phorbol ester pathway. Accumulation levels of transcripts were determined by qRT-PCR using primers in Table 1. Relative expression levels of individual transcripts in diverse tissues were calculated by comparison of field-growing mature leaf, including root, young stem, bud, young seed, yellow seed, green fruit, seedling leaf. (B) Scatter plot of the scores of the first two principal eigengene vectors derived from the principal component analysis (PCA) on the expression of genes. (C) Dendrogram plot of the PCA scores visualized the cluster of *Jatropha curcas* tissues based on the accumulation of terpene transcripts.

To obtain a systematic view of the relationships in terpenoids biosynthesis of different tissues, we conducted principal component analysis (PCA) on the terpenoid gene expression profile. The first and second principal components captured 53.68% and 42.45% of the total variance, respectively (Figure 3B). The scatter plot on the scores of PC1 and PC2 showed the accumulation patter of terpenoids transcripts varied across diverse *J. curcas* tissues (Figure 3B). Gene expression profile in seedling leaf tissue and the flowering buds showed dramatic differences from other tissues. The dendrogram based on the first two principal components divided *J. curcas* tissues into four groups: Group I (seedling leaf), Group II (Bud), Group III (young seeds and green fruits), Group IV (yellow seeds, young stem, wild leaf, and root) (Figure 3C). Collectively, the gene expression profiles of terpenoids showed tissue-specific patterns, which were associated with tissue age (i.e., young or old tissues) and tissues function (i.e., vegetative or reproductive tissues). This tissue-specific pattern of terpenoids accumulation *in planta* may link with their ecological function to protect young and reproductive tissues against abiotic and biotic stresses.

### 2.4. Cold and drought stressores increased but salinity decreased many transcript levels of terpenoids biosynthetic genes in J. curcas

Terpenoids metabolism facilitates plant adaptation to fluctuated environment and defense against biotic stresses (Pichersky and Raguso, 2018; Tholl, 2015). To examine how *Euphorbiaceae* and *Thymelaeaceae* plants coordinate terpenoid production to adapt to environmental challenges, we measured the variation of terpenoid transcripts’ accumulation in *J. curcas* seedlings upon abiotic stresses, such as cold, drought, and salinity. Besides the above eight genes involved in terpenoid common and branch pathways, we also included two genes, *SQS* and *SE*, involved in the triterpenoid pathway (Table 1 and Figure 4).

**Figure 4.**
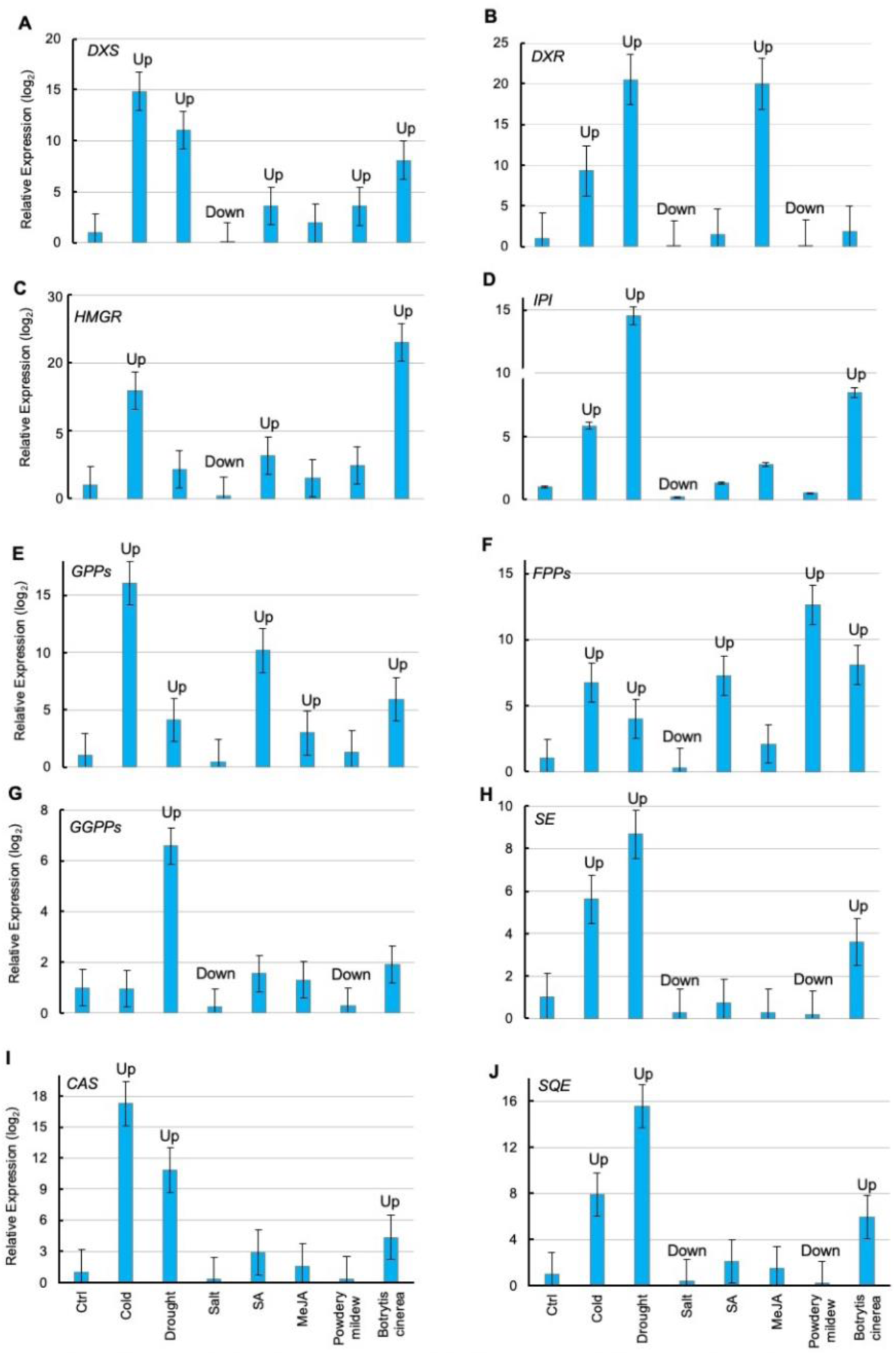
Accumulation of transcripts involved in terpene metabolism and phorbol esters pathway in response to abiotic and biotic stressors. Abiotic stressors are cold, drought, salt. Biotic stressors are plant defense hormones of salicylic acid (SA) and methyl jasmonate (MeJA), as well as biotrophic fungi *powdery mildew*, and necrotrophic fungi *Botrytis cinerea*. Accumulation levels of transcripts were determined by qRT-PCR using primers in Table 1. Relative expression levels of individual transcripts were calculated by comparison of control samples. All graphs represent the values of three replicates with cDNA prepared from seedling leaves of control samples or diverse stress treatments, respectively. Each bar represents means ± standard error.

Terpenoid gene expressions were differentially regulated by three abiotic stressors of cold, drought, and salinity (Figure 4). Cold treatment significantly affected the accumulation of terpenoids transcripts in plant tissue. When treating with cold at 4 °C for eight hours, all tested terpenoids genes showed highly expressed ranging from 5.8 to 34.8 fold change except *GGPPs* (Figure 4). In particular, *DXS* and *GPPs* showed the highest upregulation of 34.8 and 26.1 fold change after treatment, respectively (Figure 4A and 4E). Three genes (*CAS, SQS*, and *SE*) that are involved in phorbol esters and squalene productions also induced more than 10 fold change in response to low temperature (Figure 4H-4J). The upregulation of terpenoids gene expression in *J. curcas* suggests that the accumulation of terpenoids may protect plant seedlings against freezing damage. Drought stress also regulated transcripts’ accumulation of terpenoid genes but with a distinct upregulation pattern (Figure 4). For example, the gene encoding IPI in the shared isoprenoid pathway and the genes in diterpenoid and triterpenoid pathways (*SQS* and *SE*) showed dramatic upregulation in response to water deficit condition (Figure 4G, 4H, and 4J).

Strongly positive correlations were identified among the four genes based on their transcriptional accumulation levels in response to all abiotic and biotic stresses (Figure 5A). In contrast with cold and drought stresses, the high salt treatment downregulated all terpenoids genes (Figure 4). This result was different from previous studies that exposure to the drought and salinity shared many common physiological reactions in plants (Krasensky and Jonak, 2012; Seki et al., 2002; Xiong et al., 2002). The scatter plot of the scores of PC1 and PC2 highlighted that cold, drought, and salinity are distributed sparsely as PCA on terpenoid gene expression, indicating three abiotic stressors have differing effects on terpenoid transcript accumulation (Figure 5B). Collectively, the upregulation of terpenoid transcripts accumulation in plants may help plant seedlings adapt to low temperature and water-deficient conditions by increasing terpenoids content in plant cells.

**Figure 5.**
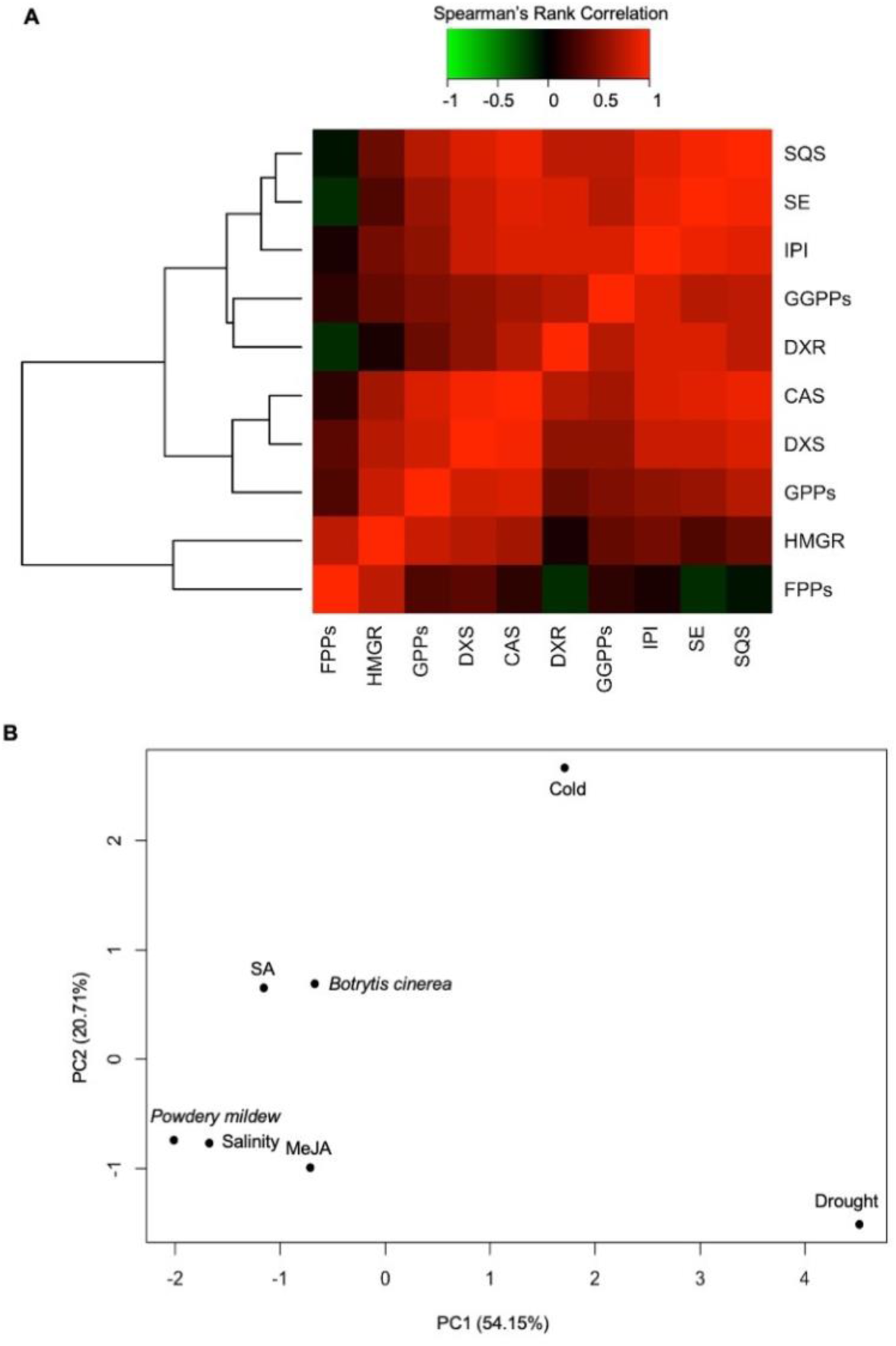
The response of genes involved in phorbol esters biosynthesis pathway to abiotic and biotic stressors. (A) Heat map illustrates the Spearman’s rank correlation co-efficiency on expression levels of target genes involved terpene metabolism and phorbol ester pathway. The accumulation levels of transcripts were determined by qRT-PCR using primers in Table 1. The cDNA was prepared from seedling leaves of control sample or under diverse abiotic and biotic stress conditions. Abiotic stressors are cold, drought, salt. Biotic stressors are plant defense hormones of salicylic acid (SA) and methyl jasmonate (MeJA), as well as biotrophic fungi *powdery mildew*, and necrotrophic fungi *B. cinerea*. Relative expression levels (log_2_) of individual transcripts were calculated by comparison with the control samples. (B) Scatter plot of the scores of the first two principal eigengene vectors derived from the principal component analysis (PCA) on the expression of the above genes. (C) Dendrogram plot of the PCA scores visualized the profile of abiotic and biotic stressors based on the terpene gene induction.

### 2.5. Attacking by biotrophic and necrotrophic fungal pathogens had different effects on transcript levels of terpenoids biosynthetic genes in J. curcas

The biotic stress effects on regulation of terpenoid gene expression were evaluated by either exogenous application of biotic hormones of salicylic acid (SA) and methyl jasmonate (MeJA) or treated with biotroph (powdery mildew, *Podosphaera xanthii*) or necrotroph (grey mode, *Botrytis cinerea*). Previous studies showed SA is essential for plant resistance upon biotrophic pathogen attack whereas JA primes plant defense against necrotrophic pathogens and insects (Kunkel and Brooks, 2002; Spoel et al., 2007; Zhang et al., 2017). Exogenous SA treatment caused the increase of all targeted terpenoids genes except *SE* (Figure 4). Specifically, the *GPPs* and *FPPs* showed the highest upregulation with 20.2 and 7.3 fold change, respectively (Figure 4E and Figure 4F). MeJA treatment showed similar but subtle induction of all targeted terpenoid genes with less than 3 fold change except *DXR*, which showed a 20 fold change (Figure 4). The varied induction patterns suggest SA and JA signaling may finely tune different terpenoids’ production.

We then investigated whether biotrophic and necrotrophic pathogen invasion trigger terpenoids gene expression by inoculation of powdery mildew (*Podosphaera xanthii*) and grey mode (*Botrytis cinerea*) on *J. curcas* seedlings (Atwell et al., 2018; Wicker et al., 2013; Zhang et al., 2016). Four genes in ten were upregulated in response to powdery mildew infection. For example, *DXS* and in shared pathway showed 3.6 and 2.4 fold change, respectively. Two genes in the monoterpenoid (*GPPs*) and sesquiterpenoid (*FPPs*) pathways showed 1.3 and 42.7 fold change, respectively (Figure 4A, 4C, 4E, and 4F). In contrast, *Botrytis cinerea* infection caused upregulation of all tested terpenoid genes and six in ten showed higher induction with more than 5 fold change upregulation compared to the control. Thus, attack by biotroph and necrotroph had different effects on terpenoid pathways at the transcriptional level in *J. curcas*. However, SA treatment and powdery mildew infection, or MeJA treatment and *Botrytis cinerea* infection were not clustered together by PCA plot (Figure 5B). Furthermore, under abiotic and biotic stresses, all tested terpenoids transcripts were positively correlated except *FPPs*. Although three clusters are identified, the genes within individual clusters are not consistent with their biosynthesis pathways, suggesting there might be crosstalks between terpenoid shared and branched pathways. Thus, the transcriptional accumulation of terpenoids is regulated *in planta* in response to diverse abiotic and biotic stresses.

## 3. Discussion

Phorbol esters are diterpenoids with broad biological activities in *J. curcas*. In this study, we determined phorbol esters distributing in whole *J. curcas* plant except for seed shells. We further showed tissue-specific patterns of the accumulation of terpenoids and phorbol esters at the transcriptional level with high transcripts levels in reproductive and young tissues. Diverse abiotic and biotic stress treatments on *J. curcas* seedling revealed that plants fine-tune gene expression in terpenoid shared and branched pathways to facilitate their fitness.

Understanding the phorbol esters distribution in organs or tissues is essential for the estimation of risks of exposure to crushed seeds or oil in agriculture and the application of *J. curcas* in medicine. This study showed phorbol esters are accumulated in all vegetative and reproductive parts in *J. curcas* but absent in seed shell (Figure 1I). Although extraction of oil can take away a majority of phorbol esters from seeds, the remained meals still contain a relatively high level of phorbol esters (Figure 1H and 1I). Thus, protection measurements are necessary to those who have occupational exposure to *J. curcas* oils and co-product of seed meals since phorbol esters are not just tumor-promoters but also skin-irritants. It is also necessary for evaluation of risks and side effects in terms of phorbol esters content when different parts of *J. curcas* plant are used in traditional medicine and pharmaceutical industry for both human and veterinary ailments.

The accumulation of phorbol esters showed a tissue-specific pattern with varied content and composition across eight tested tissues. Phorbol esters are accumulated highest in seeds, especially in endotesta, compared with leaf, stem, and latex. Further investigation of transcript accumulation revealed that genes directly responsible for phorbol esters biosynthesis are highly expressed in reproductive or young organs, including buds, green fruits, young seeds. Considering terpenoids function in defense against herbivores and pathogen attackers, phorbol esters allocation and gene expression patterns are thought to reflect the optimal allocation of defensive compounds in organs that contribute most to plant fitness. The different phorbol ester patterns in leaf, stem, flower, and seeds may be associated with specific differences of selection pressures from herbivores and pathogens acting on different plant organs. Indeed, *J. curcas* tree is also attacked and consumed by some fungal pathogens and several insects. Some of the specialists have evolved the ability to modify and store phorbol esters as their chemical defense against predators. Although the tissue-specific allocation pattern is a hallmark of most plant specialized metabolites, the driving evolutionary power remains largely unknown.

Terpenoids are the dominant secondary metabolites in *Euphorbia* plants, including *J. curcas*. We have demonstrated that the accumulation of terpenoid transcripts in *J. curcas* seedlings was upregulated in response to cold and drought stresses with a distinguished expression profile (Figure 4 and Figure 5A). Terpenoid pathways may be the crosstalks between cold and drought stressors, which were previously identified in Arabidopsis (Seki et al., 2002; Xiong et al., 2002). Further work is required to identify the molecular mechanism of the regulatory network for terpenoid production in plants in response to diverse single or combination of environmental stresses. This oilseed crop has been described as a drought-resistant perennial plant that can grow in marginal and poor soils in the tropical and subtropical regions (Reubens et al., 2011; Valdés-Rodríguez et al., 2013). Low temperature is one of the limiting factors for *J. curcas* widespread. Like many other secondary metabolites, terpenoids and the transcripts of corresponding biosynthetic genes are accumulated in response to various environmental stresses. The terpenoids production in *J. curcas* may associate with its growing environment and sever as protective metabolites that increase plants tolerance to adverse growth conditions.

As both constitutive and inducible defense compounds, terpenoids accumulation in *Euphorbia* plants may link with their ecological functions of defense against fungal pathogen and insect attack. Previous reports suggest *J. curcas* plants are suffered from several severe diseases caused by bacterial and fungal pathogens (Chowdhury et al., 2019). We showed that the expression levels of terpenoids genes are upregulated with the treatment of SA and JA or inoculation of fungi with different infection patterns (Figure 4 and Figure 5). SA and JA have been associated with the plant biotic stress hormone signals in the regulation of secondary metabolite production. Generally, SA-mediated signaling pathways are implicated in the regulation of biotrophic pathogens while the JA pathway is associated with defense responses against necrotrophic pathogens and insects (Kunkel and Brooks, 2002; Spoel et al., 2007). Terpenoids genes are regulated by SA and JA signaling pathways in distinguished patterns, suggesting different types of terpenoids production may under control by different signaling pathways. The fine-tune regulation of the specific types of terpenoid production may be associated with their ecological functions. For example, volatile terpenoids released from plants play crucial roles in attracting pollinators and repelling herbivore predators (Nagegowda, 2010). These volatile terpenoids represented by mainly isoprene, monoterpenes and sesquiterpenes may under the regulation of JA signaling pathways. Elucidating the ecological importance of terpenoids is helpful to understand and identify the molecular regulation mechanism in response to biotic stresses.

We observed that genes involved in the same type of terpenoids tends to co-expression in response to environmental signals (Figure 5A). This is not a unique feature for terpenoids production but is shared by all specialized metabolites (Schläpfer et al., 2017; Wisecaver et al., 2017). The regulation of inducible production of specialized metabolites and the transcripts involved in biosynthetic pathways in plants are coordinated at the individual gene module unit. This is the theoretical basis of the gene co-expression network approach that are broadly used in a whole-genome transcriptomic analysis to identify gene modules with associated biological functions (Soltis et al., 2020; Wisecaver et al., 2017; Zhang et al., 2017; Zhang et al., 2019). Transcriptional regulation of specialized metabolites production is under a tight regulation at different levels by many *cis*-and *trans*-regulators. Transcription factors are one of such regulators that integrate signals and interact with similar pattern of regulatory (often promoter) regions of target genes to control the specific accumulation of transcripts and specialized metabolites. Future works are required to the analysis of the patterns shared by regulatory regions of terpenoids genes and their associations with different types of transcription factors, which will help identify the underlying molecular regulatory mechanism on terpenoids induction and interpret their ecological functions.

## Conclusion

In conclusion, the accumulation of phorbol esters at metabolic and transcriptional levels in plants showed tissue-specific patterns and prefer to accumulate in reproductive and young tissues and organs. The variation of the distribution of phorbol esters across diverse tissues provides further support for the implementation of *J. curcas* plant in pharmaceutical, agricultural, and industrial applications. Furthermore, abiotic and biotic stresses had different effects on terpenoids, especially phorbol esters, accumulation in *J. curcas* seedlings at the transcriptional level. Plants can coordinate shared and branch biosynthetic pathways of terpenoids to adapt to abiotic stresses and defense against biotic stresses. This study, therefore, provides insights for understanding the ecological roles of terpenoids in plants and environment interaction.

## 4. Materials and Methods

### 4.1. Plant materials and growth condition

*J. curcas* seeds used for phorbol ester content analysis were collected in September from trees growing at the field in Panzhihua City of Sichuan Province, China. All seeds were mixed to minimize the impact of variation generated by sample differences. The kernel, shell, (episperm, the black woody part at outside of the seed), and endotesta (the white paper-like part at the outside of the kernel) were carefully separated from the above seeds. Other *J. curcas* tissues used for phorbol ester contents analysis, including root barks, stem barks, leaves, and latex were collected in July from six-year-old *J. curcas* seedlings growing in a nursery at Yanyuan County, Sichuan Province, China. The above tissues were shade dried, ground into fine powders, and stored at −20 °C until use. The latex was gathered into 1.5 mL centrifuge tubes stored at −20 °C immediately.

The *J. curcas* tissues used for gene expression profile analysis, including root, young stem, leaf, bud, young seed, yellow seed, green fruit, were collected from the same nursery and preserved in RNAlater (Ambion, Chengdu, China) at 3x the volume of RNAlater/tissue. Preserved tissue was placed at −80 °C for long-term storage. The seedling leaves were collected from the greenhouse and flash-frozen in liquid nitrogen and stored at −80 °C until RNA extraction.

### 4.2. Extraction of phorbol esters from diverse J. curcas tissues

The ultrasound-assisted extraction (UAE) method was used in this study for the extraction of phorbol esters from all types of plant material samples as described previously (Liu et al., 2013; Wang et al., 2012). *J. curcas* kernels were used in the UAE method for optimization of the five parameters, including four extraction solvents, soaking temperature, solvent/materials ratio, ultrasound irritation time, and extraction temperature. In brief, an aliquot of powdered samples was weighed into 5 mL of Eppendorf safe lock tubes. The solvent was added into samples, mixed well, and incubated for 40 min in a water bath with a fixed soaking temperature. The mixture was then sonicated in an ultrasound processor (Transsonic KH-300DB, Kunshan Hechuang, Ultrasonic Instrument Co., Ltd., Kunshan, China) for a while at a fixed temperature. The supernatant was collected. The residue remained at the bottom of the tube was re-extracted for three times using the same method. The supernatants from four extractions were pooled and evaporated for 15 min at 35 °C by a nitrogen blowing instrument to obtain an oily sample.

To minimize the impacts of coloring impurities on the phorbol esters determination from the extracts of root barks, stem barks, and leaves, we conducted a removal of pigments experiment using powdered activated carbon (PAC). The adsorption effect on phorbol esters was investigated by comparing phorbol esters’ contents between kernel extracts with and without PAC treatment. In brief, the kernel extracts were divided into two samples. One is used as a treatment by adding PAC and the other one without PAC was used as control. Samples were first dissolved in a 50-mL conical flask with methanol, only the treated sample was added with 0.2 g PAC. The samples were incubated in a water bath at 35° for 40 min to achieve a dynamic equilibrium between adsorbate and adsorbent. The mixture was filtered and filtrates were evaporated and re-dissolved in 10 mL of methanol. The Extracts A group was subjected to the same procedure as the Extracts B group to obtain a blank reading. The total phorbol esters content from two extracts were determined by HPLC.

The samples from root barks, stem barks and leaves were subjected to the same PAC treatment for removal of coloring impurities. In brief, eight aliquots of 0.300 g powder were simultaneously extracted using the optimized UAE procedure. All extracts were pooled together dissolved in methanol and subjected to PAC treatment. The PAC treatment was conducted twice. The final extracts were re-dissolved in 2 mL methanol and stored at 4 °C before HPLC analysis. The pigments present in PE extracts from root barks, stem barks, and leaves always interfere in PE detection because of the same UV absorbance at the wavelength of 280 nm as phorbol esters. We developed a purification procedure by using activated carbons as a decolorization agent. The chromatographic profile of phorbol esters from kernel extract didn’t change between purification treated samples and control samples and the yields were 5.613 mg/g ± 0.044 and 5.595 mg/g ± 0.086, respectively. The results indicated that the purification treatment showed no significant impacts on phorbol esters extraction and can be used for the removal of pigments present in colorful extracts.

### 4.3. High performance liquid chromatography

The extraction yield of phorbol esters was determined by a Waters HPLC system coupled with a 2489 UV-visible dual-wavelength absorbance detector. Phorbol esters were detected at 280 nm based on the maximum absorption wavelength measured using a TU-1900 UV-Vis spectrophotometer (Beijing Purkinje general instrument Co., Ltd., China). A Spherisorb reversed phase ODS C18 column (250 mm× 4.6 mm, 5 μm, Welch Materials, Inc., USA) was used for separating phorbol esters with the column temperature at 25 °C. The isocratic binary mobile phase consisted of ACN and water (80:20, v/v) containing 0.2% formic acid, and the flow speed was set at 1.0 mL/min. An N2000 workstation (Zhejiang University) was employed for instrument control and data acquisition. The extract was dissolved in a 10 mL amber volumetric flask in methanol and filtered before HPLC injection. The extraction and HPLC analysis were performed in triplicates. A completely randomized experimental design was used for parameter optimization. Five parameters were optimized systematically, including organic solvents, soaking temperatures, solvent-sample ratio, sonication time, and temperature.

Phorbol 12-myristate-13-acetate (PMA, CAS number 16561-29-8) was supplied by Sigma Chemical Co., Ltd. (Shanghai, China). The stock solution was prepared by dissolving the standard PMA in methanol and stored at 4 °C until use. The standard working solutions were prepared freshly by appropriate dilution of a stock solution in methanol. Seven different concentrations of working standard solutions were injected into the HPLC in triplicates. External standard PMA was present at 39 min under 280 nm detection wavelength. The peak area values were plotted versus the corresponding PMA concentrations to obtain the calibration curve. The identity of HPLC peaks in kernel extracts was based on the comparison of retention time and UV absorption spectrum as determined on a diode-array detector with those described previously (Devappa et al., 2013a; Hua et al., 2015; Liu et al., 2013; Wang et al., 2012). Phorbol esters’ yields were calculated as PMA standard equivalents and given as mg/g dry weight tissue. Total phorbol ester contents were calculated by summarizing six phorbol esters contents. The results were expressed as Means ± Standard error. The external calibration curve was plotted using the linear least-squares regression method. The linear equation was obtained: Y_PeakArea_ = 3025.3X_concentation_ −33.38 (*R*^*2*^ > 0.9998) over a linearity concentration range from 20 to 500 μg/mL. The limit of detection (LOD) and limit of quantification (LOQ) were less than 2.07 μg/mL and 6.87 μg/mL, respectively. The precision was tested by five replicated injections, and the repeatability of the peak areas was expressed as a relative standard deviation (RSD). The RSDs for PMA and PEs were 1.50% and 2.96%, respectively. Besides, peak areas and retention times of PMA and phorbol esters were found to be quite stable over 48 hours.

### 4.4. Total RNA extraction and quantitative RT-PCR analysis

Total RNA was extracted from the three independent samples from *J. curcas* seedlings. A total of 4.0 μg RNA from each sample was reverse transcribed to cDNA. Then, the three independent transcribed cDNA was used as templates for qRT-PCR. Genes involved in the phorbol ester pathway in *J. curcas* were tested and the sequences of the primers are shown in Table 1. The *J. curcas ACTIN* gene was used as a reference for relative expression calculation and ddH_2_O was used as the negative control sample. The qRT-PCR was conducted to quantify the expression levels of phorbol ester associated genes (Heid et al., 1996; Karlen et al., 2007). The procedure for qRT-PCR was as below: 1 μL of 1/10 diluted cDNA was added into 5 μL of 2x SYBR Green buffer, followed by 0.1 μM of primer pairs, and ddH_2_O to bring the volume to 10 μL.

### 4.5. Statistical analysis

The statistical calculations and analyses in this study were performed using an R programming language (R Development Core Team, 2015). Tukey’s honestly (HSD) test was conducted to identify all significant differences between means.

To test the significance of the effect of each parameter, or a specific interaction between parameters on the extraction yields, we conducted an ANOVA model containing each parameter as individual effects. The ANOVA model was phorbol esters content = solvent + soaking temperature + ratio + extraction time + extraction temperature + solvent x soaking temperature + … + soaking temperature x ratio + … + ratio x extraction time + …+ extraction time X extraction temperature. This model was used to obtain the *F*-value, significance, means, and standard error for each parameter (Supplemental Table 1).

## Author contributions and Acknowledgement

W.Z. L.W., Y.X., F.C. conceived and designed the experiments. W.Z., L.W., and S.L. performed the experiments. W.Z., L.W., T.D., and Y.X. analyzed the data. L.W., W.Z., and Y.X. wrote the paper.

## Acknowledgments

This project was supported by the National Natural Science Foundation of China Grant No. 31870315 to Y. X..

## Supplemental Data

**Supplemental Table 1.** ANOVA table for the effects of extraction parameters on yields of phorbol esters from *Jatropha curcas* kernel extracts. A linear model was used for phorbol esters yields extracted and determined by ultrasound-assisted extraction coupled with high performance liquid chromosome. Five extraction parameters were determined and compared. Df means degrees of freedom, Sum of squares means Type II Sums of Squares variation, F values means the F statistic test, *P*-value means the probability value from an F test. Significance was determined as follows: *** < 0.001; ** < 0.01, * < 0.05.

| Source of Variation | Df | Sum of squares | F value | P-Value |
| --- | --- | --- | --- | --- |
| Solvent | 3 | 1.9545 | 159.77 | < 0.001 *** |
| Soaking temperature | 6 | 0.8006 | 32.72 | 0.0079 ** |
| Ratio | 4 | 0.9491 | 58.19 | 0.0035 ** |
| Extraction time | 4 | 0.2453 | 15.04 | 0.025 * |
| Extraction temperature | 5 | 0.5936 | 29.12 | 0.009 ** |
| Total | 22 | 0.0122 |  |  |

\* $P < 0.05$ , \*\* $P < 0.01$ , \*\*\* $P < 0.001$

**supplemental Figure 1.**
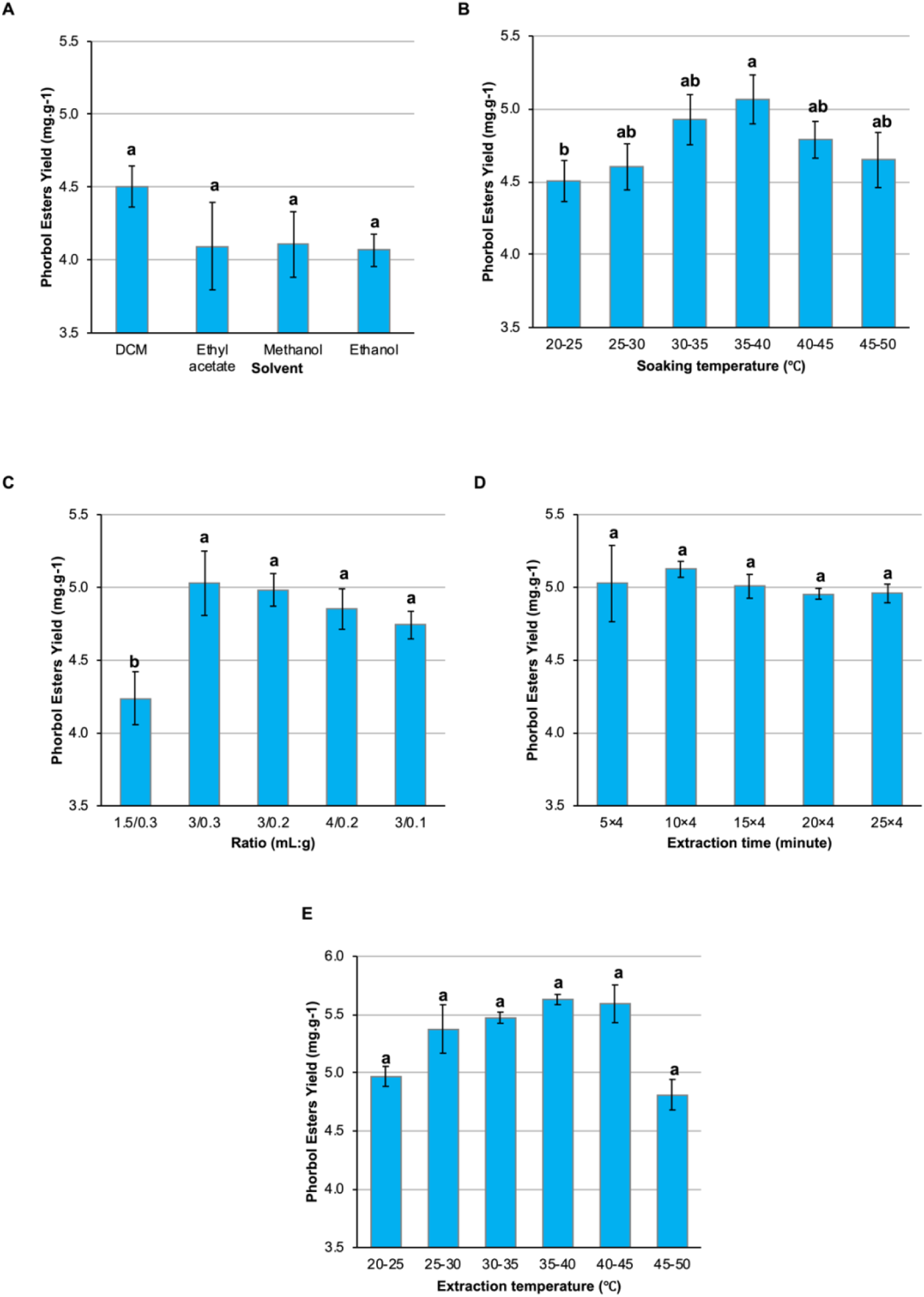
Effects of extraction parameters on the yields of phorbol esters from *Jatropha curcas* kernel extracts. (A) Effect of extraction solvents; (B) Effect of soaking temperature; (C) Effect of the solvent/material ration; (D) Effect of ultrasound irradiation time; and (E) Effect of extraction temperature. Error bars represent the standard error of the mean. Letters represent significant differences determined by Tukey’s HSD test.

## References

Atwell, S., Corwin, J.A., Soltis, N.E., Zhang, W., D., C., J., F., R., E., and J., K.D. (2018). Resequencing and association mapping of the generalist pathogen Botrytis cinerea. bioRxiv.

Becker, K., and Makkar, H. (1998). Effects of phorbol esters in carp (Cyprinus carpio L). Veterinary and Human Toxicology 40:82–86.

Berchmans, H.J., and Hirata, S. (2008). Biodiesel production from crude Jatropha curcas L. seed oil with a high content of free fatty acids. Bioresour Technol 99:1716–1721.

Chowdhury, J., Al Basir, F., Takeuchi, Y., Ghosh, M., and Roy, P. (2019). A mathematical model for pest management in Jatropha curcas with integrated pesticides - An optimal control approach.v Ecological Complexity 37:24–31.

Debnath, M., and Bisen, P.S. (2008). Jatropha curcas L., a multipurpose stress resistant plant with a potential for ethnomedicine and renewable energy. Curr Pharm Biotechnol 9:288–306.

Devappa, R., Bingham, J., and Khanal, S. (2013a). High performance liquid chromatography method for rapid quantification of phorbol esters in Jatropha curcas seed. Industrial Crops and Products 49:211–219.

Devappa, R., Maes, J., Makkar, H., De Greyt, W., and Becker, K. (2010a). Quality of Biodiesel Prepared from Phorbol Ester Extracted Jatropha curcas Oil. Journal of the American Oil Chemists Society 87:697–704.

Devappa, R., Makkar, H., and Becker, K. (2010b). Nutritional, Biochemical, and Pharmaceutical Potential of Proteins and Peptides from Jatropha: Review. Journal of Agricultural and Food Chemistry 58:6543–6555.

Devappa, R., Rajesh, S., Kumar, V., Makkar, H., and Becker, K. (2012). Activities of Jatropha curcas phorbol esters in various bioassays. Ecotoxicology and Environmental Safety 78:57–62.

Devappa, R.K., Malakar, C.C., Makkar, H.P., and Becker, K. (2013b). Pharmaceutical potential of phorbol esters from Jatropha curcas oil. Nat Prod Res 27:1459–1462.

Fujiki, H., Suttajit, M., Rawangkan, A., Iida, K., Limtrakul, P., Umsumarng, S., and Suganuma, M. (2017). Phorbol esters in seed oil of Jatropha curcas L. (saboodam in Thai) and their association with cancer prevention: from the initial investigation to the present topics. J Cancer Res Clin Oncol 143:1359–1369.

Haas, W., Sterk, H., and Mittelbach, M. (2002). Novel 12-deoxy-16-hydroxyphorbol diesters isolated from the seed oil of Jatropha curcas. Journal of Natural Products 65:1434–1440.

Heid, C.A., Stevens, J., Livak, K.J., and Williams, P.M. (1996). Real time quantitative PCR. Genome Res 6:986–994.

Hirota, M., Suttajit, M., Suguri, H., Endo, Y., Shudo, K., Wongchai, V., Hecker, E., and Fujiki, H. (1988). A new tumor promoter from the seed oil of Jatropha curcas L., an intramolecular diester of 12-deoxy-16-hydroxyphorbol. Cancer Res 48:5800–5804.

Hua, W., Hu, H., Chen, F., Tangy, L., Peng, T., and Wang, Z. (2015). Rapid Isolation and Purification of Phorbol Esters from Jatropha curcas by High-Speed Countercurrent Chromatography. Journal of Agricultural and Food Chemistry 63:2767–2772.

Igbinosa, O., Igbinosa, I., Chigor, V., Uzunuigbe, O., Oyedemi, S., Odjadjare, E., Okoh, A., and Igbinosa, E. (2011). Polyphenolic Contents and Antioxidant Potential of Stem Bark Extracts from Jatropha curcas (Linn). International Journal of Molecular Sciences 12:2958–2971.

Insanu, M., Dimaki, C., Wilkins, R., Brooker, J., van der Linde, P., and Kayser, O. (2013). Rational use of Jatropha curcas L. in food and medicine: from toxicity problems to safe applications. Phytochemistry Reviews 12:107–119.

Joseph, G.V. (2001). Pharmacognostic studies on the fruits of jatropha curcas linn. Anc Sci Life 21:128–134.

Karlen, Y., McNair, A., Perseguers, S., Mazza, C., and Mermod, N. (2007). Statistical significance of quantitative PCR. BMC Bioinformatics 8:131.

Katole, S., Saha, S., Sastry, V., Lade, M., and Prakash, B. (2011). Intake, blood metabolites and hormonal profile in sheep fed processed Jatropha (Jatropha curcas) meal. Animal Feed Science and Technology 170:21–26.

King, A., Brown, G., Gilday, A., Larson, T., and Grahama, I. (2014). Production of Bioactive Diterpenoids in the Euphorbiaceae Depends on Evolutionarily Conserved Gene Clusters. Plant Cell 26:3286–3298.

King, A.J., He, W., Cuevas, J.A., Freudenberger, M., Ramiaramanana, D., and Graham, I.A. (2009). Potential of Jatropha curcas as a source of renewable oil and animal feed. J Exp Bot 60:2897–2905.

Kosasi, S., Vandersluis, W., Boelens, R., Thart, L., and Labadie, R. (1989). LABADITIN, A NOVEL CYCLIC DECAPEPTIDE FROM THE LATEX OF JATROPHA-MULTIFIDA L (EUPHORBIACEAE). Febs Letters 256:91–96.

Krasensky, J., and Jonak, C. (2012). Drought, salt, and temperature stress-induced metabolic rearrangements and regulatory networks. J Exp Bot 63:1593–1608.

Krishnamurthy, L., Zaman-Allah, M., Marimuthu, S., Wani, S., and Rao, A. (2012). Root growth in Jatropha and its implications for drought adaptation. Biomass & Bioenergy 39:247–252.

Kunkel, B.N., and Brooks, D.M. (2002). Cross talk between signaling pathways in pathogen defense. Curr Opin Plant Biol 5:325–331.

Lange, B.M., Rujan, T., Martin, W., and Croteau, R. (2000). Isoprenoid biosynthesis: the evolution of two ancient and distinct pathways across genomes. Proc Natl Acad Sci U S A 97:13172–13177.

Li, C., Devappa, R., Liu, J., Lv, J., Makkar, H., and Becker, K. (2010). Toxicity of Jatropha curcas phorbol esters in mice. Food and Chemical Toxicology 48:620–625.

Li, C., Ng, A., Xie, L., Mao, H., Qiu, C., Srinivasan, R., Yin, Z., and Hong, Y. (2016). Engineering low phorbol ester Jatropha curcas seed by intercepting casbene biosynthesis. Plant Cell Reports 35:103–114.

Lin, J., Li, Y.X., Zhou, X.W., Tang, K.X., and Chen, F. (2003). Cloning and characterization of a curcin gene encoding a ribosome inactivating protein from Jatropha curcas. DNA Seq 14:311–317.

Liu, X., Li, L., Li, W., Lu, D., Chen, F., and Li, J. (2013). Quantitative determination of phorbol ester derivatives in Chinese Jatropha curcas seeds by high-performance liquid chromatography/mass spectrometry. Industrial Crops and Products 47:29–32.

Makkar, H., Becker, K., Sporer, F., and Wink, M. (1997). Studies on nutritive potential and toxic constituents of different provenances of Jatropha curcas. Journal of Agricultural and Food Chemistry 45:3152–3157.

Makkar, H., Francis, G., and Becker, K. (2008). Protein concentrate from Jatropha curcas screw-pressed seed cake and toxic and antinutritional factors in protein concentrate. Journal of the Science of Food and Agriculture 88:1542–1548.

Montes, J.M., and Melchinger, A.E. (2016). Domestication and Breeding of Jatropha curcas L. Trends Plant Sci 21:1045–1057.

Nagegowda, D.A. (2010). Plant volatile terpenoid metabolism: biosynthetic genes, transcriptional regulation and subcellular compartmentation. FEBS Lett 584:2965–2973.

Nakano, Y., Ohtani, M., Polsri, W., Usami, T., Sambongi, K., and Demura, T. (2012). Characterization of the casbene synthase homolog from Jatropha (Jatropha curcas L.). Plant Biotechnology 29:185–189.

Pichersky, E., and Raguso, R.A. (2018). Why do plants produce so many terpenoid compounds? New Phytol 220:692–702.

Qin, W., Ming-Xing, H., Ying, X., Xin-Shen, Z., and Fang, C. (2005). Expression of a ribosome inactivating protein (curcin 2) in Jatropha curcas is induced by stress. J Biosci 30:351–357.

Ravindranath, N., Ravinder Reddy, M., Ramesh, C., Ramu, R., Prabhakar, A., Jagadeesh, B., and Das, B. (2004). New lathyrane and podocarpane diterpenoids from Jatropha curcas. Chem Pharm Bull (Tokyo) 52:608–611.

Reubens, B., Achten, W., Maes, W., Danjon, F., Aerts, R., Poesen, J., and Muys, B. (2011). More than biofuel? Jatropha curcas root system symmetry and potential for soil erosion control. Journal of Arid Environments 75:201–205.

Rug, M., and Ruppel, A. (2000). Toxic activities of the plant Jatropha curcas against intermediate snail hosts and larvae of schistosomes. Tropical Medicine & International Health 5:423–430.

Sabandar, C., Ahmat, N., Jaafar, F., and Sahidin, I. (2013). Medicinal property, phytochemistry and pharmacology of several Jatropha species (Euphorbiaceae): A review. Phytochemistry 85:7–29.

Saetae, D., and Suntornsuk, W. (2011). Toxic Compound, Anti-Nutritional Factors and Functional Properties of Protein Isolated from Detoxified Jatropha curcas Seed Cake. International Journal of Molecular Sciences 12:66–77.

Schläpfer, P., Zhang, P., Wang, C., Kim, T., Banf, M., Chae, L., Dreher, K., Chavali, A.K., Nilo-Poyanco, R., Bernard, T., et al. (2017). Genome-Wide Prediction of Metabolic Enzymes, Pathways, and Gene Clusters in Plants. Plant Physiol 173:2041–2059.

Seki, M., Narusaka, M., Ishida, J., Nanjo, T., Fujita, M., Oono, Y., Kamiya, A., Nakajima, M., Enju, A., Sakurai, T., et al. (2002). Monitoring the expression profiles of 7000 Arabidopsis genes under drought, cold and high-salinity stresses using a full-length cDNA microarray. Plant J 31:279–292.

Soltis, N.E., Caseys, C., Zhang, W., Corwin, J.A., Atwell, S., and Kliebenstein, D.J. (2020). Pathogen Genetic Control of Transcriptome Variation in the. Genetics 215:253–266.

Spoel, S.H., Johnson, J.S., and Dong, X. (2007). Regulation of tradeoffs between plant defenses against pathogens with different lifestyles. Proc Natl Acad Sci U S A 104:18842–18847.

Tholl, D. (2015). Biosynthesis and biological functions of terpenoids in plants. Adv Biochem Eng Biotechnol 148:63–106.

Valdés-Rodríguez, O.A., Sánchez-Sánchez, O., Pérez-Vázquez, A., Caplan, J.S., and Danjon, F. (2013). Jatropha curcas L. root structure and growth in diverse soils. ScientificWorldJournal 2013:827295.

Vranová, E., Coman, D., and Gruissem, W. (2013). Network analysis of the MVA and MEP pathways for isoprenoid synthesis. Annu Rev Plant Biol 64:665–700.

Wang, Z., Tang, L., Hu, H., Guo, Y., Peng, T., Yan, F., and Chen, F. (2012). Metabolic Profiling Assisted Quality Control of Phorbolesters in Jatropha curcas Seed by High-Performance Liquid Chromatography Using a Fused-Core Column. Journal of Agricultural and Food Chemistry 60:9567–9572.

Wei, L., Zhang, W., Yin, L., Yan, F., Xu, Y., and Chen, F. (2015). Extraction optimization of total triterpenoids from Jatropha curcas leaves using response surface methodology and evaluations of their antimicrobial and antioxidant capacities. Electronic Journal of Biotechnology 18:88–95.

Wicker, T., Oberhaensli, S., Parlange, F., Buchmann, J.P., Shatalina, M., Roffler, S., Ben-David, R., Doležel, J., Šimková, H., Schulze-Lefert, P., et al. (2013). The wheat powdery mildew genome shows the unique evolution of an obligate biotroph. Nat Genet 45:1092–1096.

Wisecaver, J.H., Borowsky, A.T., Tzin, V., Jander, G., Kliebenstein, D.J., and Rokas, A. (2017). A Global Coexpression Network Approach for Connecting Genes to Specialized Metabolic Pathways in Plants. Plant Cell 29:944–959.

Xiong, L., Schumaker, K.S., and Zhu, J.K. (2002). Cell signaling during cold, drought, and salt stress. Plant Cell 14 Suppl:S165–183.

Yazaki, K., Arimura, G.I., and Ohnishi, T. (2017). ‘Hidden’ Terpenoids in Plants: Their Biosynthesis, Localization and Ecological Roles. Plant Cell Physiol 58:1615–1621.

Zerbe, P., and Bohlmann, J. (2015). Plant diterpene synthases: exploring modularity and metabolic diversity for bioengineering. Trends Biotechnol 33:419–428.

Zhang, W., Corwin, J.A., Copeland, D., Feusier, J., Eshbaugh, R., Chen, F., Atwell, S., and Kliebenstein, D.J. (2017). Plastic Transcriptomes Stabilize Immunity to Pathogen Diversity: The Jasmonic Acid and Salicylic Acid Networks within the Arabidopsis/Botrytis Pathosystem. Plant Cell 29:2727–2752.

Zhang, W., Corwin, J.A., Copeland, D., Feusier, J., Eshbaugh, R., Cook, D.E., Atwell, S., and Kliebenstein, D.J. (2019). Plant-necrotroph co-transcriptome networks illuminate a metabolic battlefield. Elife 8.

Zhang, W., Kwon, S.T., Chen, F., and Kliebenstein, D.J. (2016). Isolate Dependency of Brassica rapa Resistance QTLs to Botrytis cinerea. Frontiers in Plant Science 7.

